# Ability of the Ash dieback pathogen to reproduce and to induce damage on its host are controlled by different environmental parameters

**DOI:** 10.1101/2022.05.03.490390

**Authors:** Benoit Marçais, Arnaud Giraudel, Claude Husson

**Affiliations:** Université de Lorraine, INRAE-Grand-Est, UMR1136 Interactions Arbres/Microorrganismes, F-54000 Nancy, France; Département de la Santé des Forêts, Ministère de l’Agriculture et de l’Alimentation - DGAL, F-75015 Paris, France

**Keywords:** *Hymenoscyphus fraxineus*, climate, healthy carrier, latent infection, apothecia production

## Abstract

Ash dieback, induced by an invasive ascomycete, *Hymenoscyphus fraxineus*, has emerged in the last decade as a severe disease threatening ash populations in Europe. Future prospects for Ash are improved by the existence of individuals with natural genetic resistance to the disease and by limited disease impact in many environmental conditions where ashes are frequent. Nevertheless, it was suggested that even in those conditions, ash trees are infected and enable pathogen transmission. We studied the influence of climate and local environment on the ability of *H. fraxineus* to infect, be transmitted and cause damage on its host. We showed that healthy carrier, i.e. asymptomatic individuals carrying *H. fraxineus*, exists and may play a significant role in ash dieback epidemiology. Environment strongly influenced *H. fraxineus* with different parameters being important depending on the life cycle stage. The ability of *H. fraxineus* to establish on ash leaves and to reproduce on the leaf debris in the litter (rachises) mainly depended on total precipitations in July-August and was not influenced by local tree cover. By contrast, damages to the host, and in particular shoot mortality was significantly reduced by high summer temperature in July-August and by high autumn average temperature. As a consequence, in many situations ash trees are infected and enable *H. fraxineus* transmission while showing limited or even no damages. We also observed a decreasing trend of severity (leaf necrosis and shoot mortality likelihood) with the time of disease presence in a plot that could be significant for the future of Ash dieback.

## Introduction

The existence of heathy carrier, i.e. infected individual that remain asymptomatic, is not recognized as an important feature in plant epidemiology (Cunniffe et al, 2015). The existence of heathy carriers have especially been reported for weak pathogens or those that can induce damage only on stressed hosts (so-called latent pathogens, Sieber, 2007). This is known to be a significant feature for important stress-induced trees diseases such as *Diplodia* shoot blight, sooty bark disease of maples or *Botryosphaeria* cankers (Flowers et al, 2001, Marsberg et al, 2017, Kelnarová et al, 2017). However, high production of inoculum usually occurs only at symptoms onset. By contrast, existence of heathy carriers is seldom reported for primary pathogens that do not need host stress to induce significant host damages. Tolerance has been reported as a mechanism used by plant to cope with infection (Pagán & García-Arenal, 2020). This mechanism postulate that some individuals will limit the impact of infection on their fitness while sustaining significant pathogen load and enabling pathogen multiplication. However, the presence of individual plants that provide important levels of inoculum while exhibiting limited or even no visible symptoms is seldom reported. Disease severity is usually assumed to correlate positively with the multiplication of the pathogen within the host (Alizon & Michalakis, 2015). Nevertheless, at the inter-species level, it has been shown that poor correlation may exist between vulnerability (reduction of fitness caused by the infection) and competence (ability to sustain the epidemic by providing large amount of inoculum). A striking example is given by the Sudden Oak Death that developed in California in the last decades: while this disease, induced by *Phytophthora ramorum*, affects primarily *Quercus* and *Lithocarpus* species, the main inoculum producer is a laurel, *Umbellularia californica* which coexist in natural forest (so-called reservoir host, Davidson et al, 2005, Garbelotto et al, 2017).

Such a discrepancy between pathogen’s ability to establish in a stand and produce inoculum and it ability to induce damages is an important difficulty for assessing the climatic niche of a pathogen. This assessment is a critical step to evaluate the risk associated with pathogen occurrence (Holdenrieder et al, 2004). The most widespread approach to develop species distribution models (SDM) is to relate the species known occurrence to environmental parameters, in particular climate, through statistical analyses (Elith & Leathwick, 2009). If large discrepancies exist between a pathogen presence and its ability to cause serious damage, this will lead to important under-reporting and to poor available data. By contrast, mechanistic SDM, which relies on the assessment of environment influence on the different stages of the pathogen cycle (Dormann et al, 2012) could be a suitable alternative to assess the climatic niche of a pathogen in such conditions.

*Hymenoscyphus fraxineus* is an invasive pathogen that has caused Ash dieback throughout Europe since the late nineties and has jeopardized *Fraxinus excelsior* stands (Gross et al, 2014). *H. fraxineus* fulfils its life cycle on ash leaves. Those are infected by ascospores during late spring and summer and remain asymptomatic until late August (Cross et al, 2017). *H. fraxineus* then colonizes the leaf petiole and may infect young shoots, inducing dieback (Gross et al, 2014). Most of the inoculum production occurs on leaf debris in the forest litter (so-called rachis that encompasses the petiole and the main vein of the composite leave, on which fruit bodies i.e. apothecia develop, Gross et al, 2014). By contrast, very limited inoculum production occurs on diseased shoots which is considered to represent a dead-end for *H. fraxineus* (Landolt *et al*, 2016). It has been suggested that, in some conditions, *H. fraxineus* may reproduce on Ash leaves without inducing significant dieback. First, it has been shown that some *F. excelsior* individuals are not affected by *H. fraxineus*, even under high inoculum pressure (McKinney et al, 2011). These individuals did not sustain extensive lesions when artificially inoculated at the shoot level (McKinney et al, 2012). It was then suggested that their leaves can nevertheless sustain the pathogen reproduction and that the mechanism involved in containing the disease is tolerance instead of resistance (Landolt *et al*, 2016). However, no experimental data is yet available to sustain this claim. Second, Grosdidier et al (2020) showed that in open canopy conditions, the infection levels of ash rachis by *H. fraxineus* in the litter was similar compared to forest stands with a close canopy despite much lower level of crown dieback. Those data would suggest that healthy carriers could exist in the *F. excelsior* population, with individuals able to be infected in specific conditions at the leaf level and thus allow to produce inoculum while sustaining very limited to no dieback.

We thus examined whether Ash trees could in some conditions behave as healthy carriers. More specifically, we studied environmental influence on important stages of Ash dieback, namely the ability to colonize leaf rachises and reproduce on them and the ability to induce damage on ash trees, either leaf necrosis at the end of the summer or subsequently shoots mortality. The aim was to provide basis for a latter development of a mechanistic SDM by documenting the key steps of *H. fraxineus* life cycle.

## Material and methods

### Sampled plots

We selected plots on a temperature gradient in France (Fig. 1) in order to have a range of summer temperature as large as possible within the range *of H. fraxineus* presence. The plots were usually studied several years in a row starting from 2013 in area affected by ash dieback since 2009-10 (NE France) to 2020 in an area just recently affected by Ash dieback (Brittany, close to Brest and Pyrenean Piedmont, close to Pau). The studied plots were either in forest settings or located in hedges. The ash species present was *Fraxinus excelsior* in all plots. A summary of observation of leaf necrosis and shoot mortality on the plots is given in table S1. Plots were characterized by their tree cover from aerial photographs (Orthophoto with 50cm pixels from IGN, http://professionnels.ign.fr). The tree cover polygon within a radius of 100 m from each plot were extracted and the proportion of tree cover within that range (TC) was computed.

**Fig. 1.**
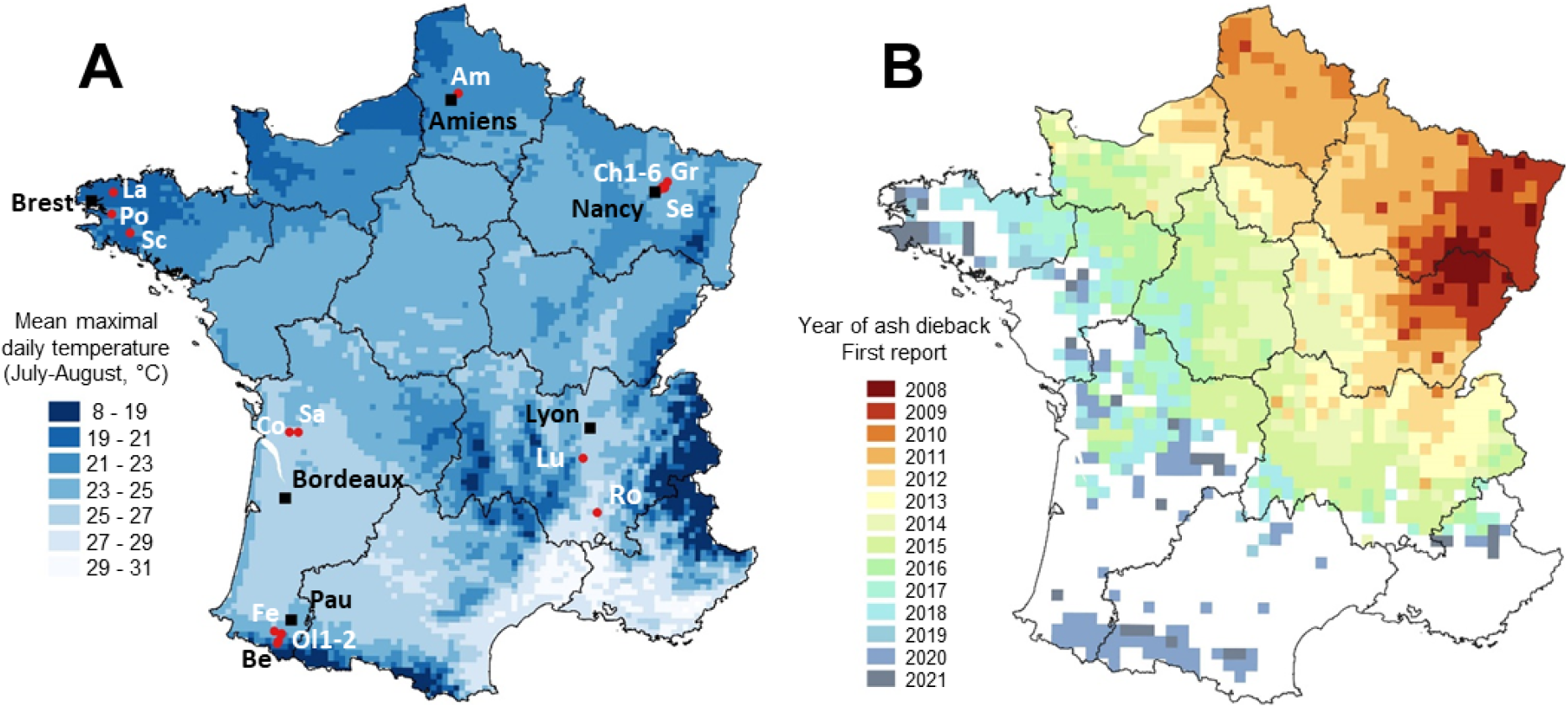
Location of the studied ash stands (red dot) relative to climate (A) and year of first ash dieback report (B, data from the Département de la Santé des Forêts). Am, Fréchencourt, Be, Sarrance, Ch1-6, Champenoux (6 plots), Co, Colombier, Fe, Ance-Féas, Gr, Gremeçey, La, Landivisiau Lu, Lupé, Ol1-2, Oloron-Sainte-Marie (2 plots), Po, Pont-de-Buis, Ro, Roche-sur-Grane, Sa, Salignac-sur-Charente Sc, Scaёr, Se, Seichamps)

### Survey procedure

In all studied plots, 20 ashes were randomly selected. We usually selected sapling 2-4m high, although occasionally, larger trees with low lying branches were selected. The saplings were rated for their decline status according to five dieback scores: 0 for complete absence of dead twigs, 1 for less than 10% of dead twigs, 2 for 10 to 50% of dead twigs, 3 for more than 50 % of dead twigs. Two annual shoots per individual were marked for rating of the leaf infection in autumn. We selected the terminal shoot of the 2 lower branches that could be reached by an observer from the ground.

The necrosis induced by *H. fraxineus* on leaves was rated in September. The total number of leaves present, the number of leaves with lesions and the number of leaf scars indicating defoliation were determined. Lesions induced by *H. fraxineus* can be recognized as brown lesions starting at leaf margin and extending preferentially on leaf veins. Leaves presenting brown lesions only on the rachis were also rated as infected. Two leaves rated as presenting *H. fraxineus-induced* lesions were sampled for further analysis in the laboratory in 2018-20. Sampled leaves were analyzed by qPCR for the presence of *H. fraxineus* as described by Ioos et al. (2009). In the following year vegetation season, the status of marked shoots was assessed as living or dead. Whenever the shoot had wilted because of girdling occurring below the annual shoot, the data was counted as missing. Saplings were rated during 1-6 years depending on the sites (Table S1). Dead twigs were replaced by the nearest shoot available from the lower part of the sapling. Dead saplings were replaced taking care to select saplings with ash dieback symptoms to avoid ending with only individuals potentially tolerant to *H. fraxineus*.

In October-November, 20 rachises that had just fallen to the ground were sampled by site in order to assess the ability of *H. fraxineus* to produced apothecia on them (called hereafter fructification assay). This was done in all sites of Champenoux (2016, 2017, 2019, 2020), Fréchencourt (2016, 2017), Lupé (2016), Roche-sur-Grane (2017) and in the 4 sites close to Pau (2020). The rachises were placed in mesh bags and overwintered in common garden in a forest site in Champenoux (`6.34251, 48.75297`, EPSG 4326) except for the rachises of the plots close to Pau that were overwintered close to the site of Sarrance (−0.59692, 43.00044, EPSG 4326). In the following spring, in April or May depending on the year, the rachises were retrieved from the mesh bags and were placed for 6-8 weeks at 18-20°C in humid chamber in large plastics boxes in contact with moistened filter paper to assess their ability to produce apothecia. The moistened filter paper was kept wet throughout the incubation and was changed weekly. Whenever a rachis produced *H. fraxineus* apothecia, it was removed and stored in paper bags for further analysis. *H. fraxineus* apothecia can be recognized as 2-4 mm white apothecia with a dark base of the pedicels and smooth margin of the fruiting disk (Kowalski & Holdenrieder, 2009). The monitoring was stopped whenever no new production of apothecia was observed in a week (after 6-8 weeks incubation). Remaining rachises were then placed in a separate bags. The amount of rachises producing / not producing apothecia was determined by measuring the rachis length in 2016 and 2017 and by measuring their weight after drying 3d at 50°C in 2019 and 2020. The proportion of rachises producing apothecia was then computed.

In 2019 and 2020 in sites of Champenoux and in 2020 in sites close to Pau, 10 additional rachises were collected per site at the same time at leaf fall to determine the frequency of *H. fraxineus* presence by qPCR. Individual rachises were kept separate. They were dried 3d at 50°C, cut in small 1-2 mm sections and grinded with a mortar. A subsample of 50 mg per rachis was the used for a DNA extraction with DNAeasy Plant Mini kit (QIAGEN). AP1 lysis buffer (800 μl), 4 μl RNase, two 3-mm and twenty 2-mm glass beads were added to the tube and the sample was ground twice using a beadbeater (Tissue Lyser,Qiagen) at 30 Hz for 2× 30 s. The DNA extraction was then conducted according to the manufacturer’s recommendations. Total DNA was eluted in 150 μl AE buffer. The samples were then analysed by qPCR for the presence of *H. fraxineus* as described in Ioos et al. (2009).

### Ability of *H. fraxineus* to complete its cycle on asymptomatic and severely affected ash saplings

The ability of *H. fraxineus* to complete it cycle on asymptomatic ash saplings was studied in two of the plots, Gremecey and Seichamps (naturally regenerated ash stands). Ash saplings about 2-3 m high were selected during summer 2014 in order to have 10-13 saplings of two types in each stand: i. asymptomatic saplings with either no or only very few scattered small dead shoots, ii. saplings strongly affected, with top shoots killed by *H. fraxineus*. The two stands were densely stocked stands heavily infected by *H. fraxineus* since at least 4 years. The asymptomatic saplings were in close vicinity to heavily declining individuals, with usually several severely affected saplings within less than one meter from the asymptomatic sapling.

On each sapling, an annual shoot with 7-28 leaves was selected and marked at the end of July. The selected shoot infection by *H. fraxineus* was rated at the beginning of September with record of the number of leaves with necrosis and total number of leaves. To measure the ability of *H. fraxineus* to complete its cycle on rachises, we placed a net around the entire crown of each sapling in mid-September to collect all the leaves falling during the autumn. The nets containing the leaves were retrieved at the end of October and placed on the ground outdoor in a nursery close to both plots to overwinter. Unfortunately, management occurred in the Seichamps plot during the autumn and left only 4 asymptomatic and 1 declining saplings available in that plot for the net harvest. In April 2016, the rachises were retrieved from the 27 nets (11+4 asymptomatic and 11+1 severely affected by *H. fraxineus*) and were placed in large plastics boxes in contact with moistened filter paper. The boxes were then closed and placed in the laboratory at room temperature. After a 5-week incubation, the rachises were examined for presence of *H. fraxineus* apothecia. The total length of each rachis was determined as well as the length between the 2 most distant apothecia of the rachis (hereafter called L.apo). The percentage of rachis colonization on a sapling was determined as ∑ L.apo / ∑ total rachis length. The saplings were again assessed in summer 2015 to determine the global infection status and the presence of shoot infection on the marked shoot assessed in 2014 for leaf infection.

The likelihood of leaf necrosis in September 2014 and shoot infection in summer were assessed by logistic regression with the plot, the health status of the saplings and their interaction as independent variables. We used a quasibinomial distribution to analyze leaf necrosis likelihood because data showed over dispersion. The percent rachis colonization by *H. fraxineus* in spring 2015 was analyzed by beta regression using the betareg package. The stand*health status interaction was not assessed for shoot infection and % rachis colonization in 2015 as very few saplings remained in Seichamps

### Data analysis

During survey, we noticed that some of the rated shoots were very vigorous with many rated leaves and that they were seldom infected by *H. fraxineus* whatever the general situation in the stand. Preliminary analysis showed that shoots with more than 20 leaves had on average 5.9% necrotic leaves and no shoot mortality, compared to average values of 16.0 % of necrotic leaves and 21.6% shoot mortality for the total dataset. Shoots with more than 20 leaves were thus removed from the data set prior analysis. Two different models were developed.

#### Relationship between leaf necrosis and shoot mortality

The first model aimed at exploring the relationship between leaf necrosis and shoot mortality at the individual shoot level and how this may be controlled by tree health and site tree cover. Data were thus kept at the shoot level for this model. A hierarchical Bayesian model was developed:

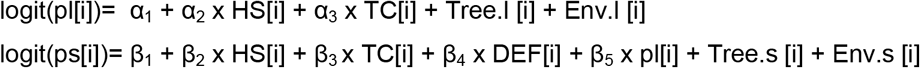

with pl[i] and ps[i] the respective probability of leaf necrosis and shoot mortality, HS[i] the ash health status (either healthy, when crown rating was of 0 or 1 or declining, when crown status was of 2 or 3), TC[i] the percent tree cover within a radius of 100 m, DEF[i] the proportion of missing leaves at the autumn rating, Tree.l [i] and Tree.s [i] individual tree random factors and Env.l [i] and Env.s [i], plot[i]*year[i] random factors. The Env.l [i] and Env.s [i] random factors were taken as proxies for the environment, combining meso/micro climate and plot conditions (host density, Infection history). The defoliation level (DEF[i]) was assumed to influence the shoot mortality likelihood because McKinney & al (2011) suggested that early defoliation may prevent the transfer of *H. fraxineus* from leaves to shoots and thus represent a tolerance mechanism.

Both pl[i] and ps[i] were assumed to follow binomial distribution. Non-informative priors were assigned to the model parameters according to a normal distribution N(0, 1e+05) or a uniform distribution U(0,100) for the standard deviation of the random factors Tree and Env. The model was fitted with a burn-in of 3000, for 50000 iterations with a thinning of 5. Three parallels chains with different initial parameter values were run and convergence was checked by a Gelman-Rubin test (R_hat_ were close to 1). The model was checked by looking at the plot of deviance residuals versus predicted values. The model was fitted with Jags 4.3.0 emulated with R4.1.0 (R2jags library).

#### Relationship between leaf necrosis or shoot mortality and climate

The second developed hierarchical Bayesian model aimed at exploring the relationship between leaf necrosis or shoot mortality with climate. For that model, data were pooled at the plot * year level. As in the previous model, shoot mortality was assumed to derive from former leaf infection and the proxy for leaf infection was taken as the leaf necrosis probability. Length of ash dieback presence in the area was included in the model to explain the likelihood of leaf necrosis. The year of first report of Ash dieback by the health survey system is available by 16 × 16 km quadrat over France from data of the health survey system (Département de la Santé des Forêts, Fig. 1). The number of years since the report of Ash dieback in the local quadrat was taken as a proxy for the length of the disease presence. It was considered an adequate proxy as local spread of *H. fraxineus* at the scale of a village was shown to be very fast (Grosdidier & al, 2020). Leaf necrosis likelihood was assumed to depend on spring and summer climatic conditions (sum of rain and average temperatures). It is known that *H. fraxineus* may not survive well during hot summers (Hauptmann & al, 2013). In particular, this pathogen is known to stop his growth at temperature above 28°C and to have limited survival when temperature exceeds 36°C. To account for this, we used TX78, the average daily maximal temperature in July-August (°C). Shoot mortality was assumed to depend on summer and autumn climatic conditions (sum of rain and average temperatures). The meteorological data used were Safran data from Météo-France. Safran data are computed on a 8 × 8 km grid over France (Quintana Segui & al, 2008). The Safran point the closest to each site was selected and daily air temperature and rain data were retrieved for the studied years.

In a first step, the suitable periods for spring and autumn were selected by a preliminary analysis. For leaf necrosis a beta regression with a logit link function was used, as the number of observed leaves per site was very high (from 200 to 400). For shoot mortality, a logistic regression with a logit link was used. The site was included as a random factor in both models as well as the frequency of leaf necrosis in the site*year for shoot mortality. The model were fitted using the glmmTMB library. The explored climate variables targeted either the ascospore production in late spring or the development of shoot necrosis during the autumn. As the adequate period for ascospore production or for the development of shoot necrosis is not known precisely, several periods were tested: from 15 May to 15 June, June or from the 15 June to 15 July for spring and September to October or September to December for autumn. For both models, climatic parameters included mean temperature and sum of rain. All predictors were normalised to obtain zero mean and one standard deviation.

In a second step, the hierarchical Bayesian model was fitted (Figueroa-Zúñiga & al, 2013):

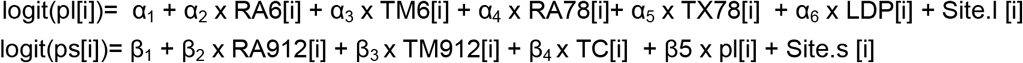

with pl[i] and ps[i] the respective likelihood of leaf necrosis and shoot mortality, RA6[i] and TM6[i] the sum of rain and mean daily air temperature in June, RA78[i] and TX78[i] the sum of rain and average daily air maximal temperature in July-August (°C), LDP[i] the number of year of ash dieback presence in the area, RA912[i] and TM912[i], the sum of rain and mean daily air temperature in September, October, November and December and TC[i] the percent tree cover within a radius of 100m. Site.l [i] and Site.s [i] are site random factors, taken as a proxy for the site environment (micro climate, site conditions such as host density or Infection history).

While ps[i] were assumed to follow binomial distribution, pl[i] was assumed to follow a beta distribution. Non-informative priors were assigned to the model parameters according to a normal distribution N(0, 1e+05) or a uniform distribution U(0,100) for the standard deviation of the random factors. The model was fitted with a burn-in of 10000, for 80000 iterations with a thinning of 10. Three parallels chains with different initial parameter values were run and convergence was checked by a Gelman-Rubin test (R_hat_ were close to 1). The model was checked by looking at the plot of deviance residuals versus predicted values and by a cross validation procedure. Data were randomly separated in 5 equal groups and the model was fitted using the data from 4 of the groups; predicted leaf necrosis frequency and shoot mortality were computed for the fifth group not used for the fit. The procedure was repeated five time and the observed value obtained by the procedure were plotted against the predicted values.

#### Relationship between climatic parameters and ability of *H. fraxineus* to produce apothecia in the fructification assay

The proportion of rachises producing apothecia in the fructification assay was analyzed by beta regression using the glmmTMB library. The site was included as random factor. Fixed factors were the proportion of leaves with necrosis at the September rating, climatic parameters in the studied sites during the summer (sum of rain and mean of the daily maximal temperature in July and August) and climatic parameters during the overwintering period (average temperature and sum of rain at the overwintering locations from beginning of October to end of March).

## Results

Altogether, data were available for 55 plot*year combinations for leaf necrosis frequency. Less data was available for shoot mortality (44 site*year combinations) because some plots were lost during winter (logging, 4 plots) or were not rated in spring (Table S1). Altogether, *H. fraxineus* was detected by the qPCR assay in 86% of the 63 tested putatively infected leaves. However the result depended on the plot and year, with lower detection frequency in plots with very few leaf necrosis (75% of tested putatively infected leaves when leaf necrosis frequency was lower than 5%; 84% when it was between 5 and 15%; and 95% when it was higher than 15%). So, assessment of leaf necrosis frequency is correct when leaf necrosis is very frequent, but may be slightly overestimated when very low.

### Ability of *H. fraxineus* to complete its cycle on ash saplings with different dieback severity

The percent of leaves with necrosis in early September 2014 was similar on asymptomatic and severely affected saplings (likelihood Chi-square of 0.7084, p= 0.400, Fig 2a). The likelihood of leaf necrosis was significantly higher in Gremecey than in Seichamps (likelihood Chi-square= 4.55, p= 0.033, 38.0 ± 6.5 % in Gremecey and 20.6 ± 4.4 % in Seichamps) but the plot * health status interaction was not significant (likelihood Chi-square = 1.32, p= 0.251).

**Fig. 2.**
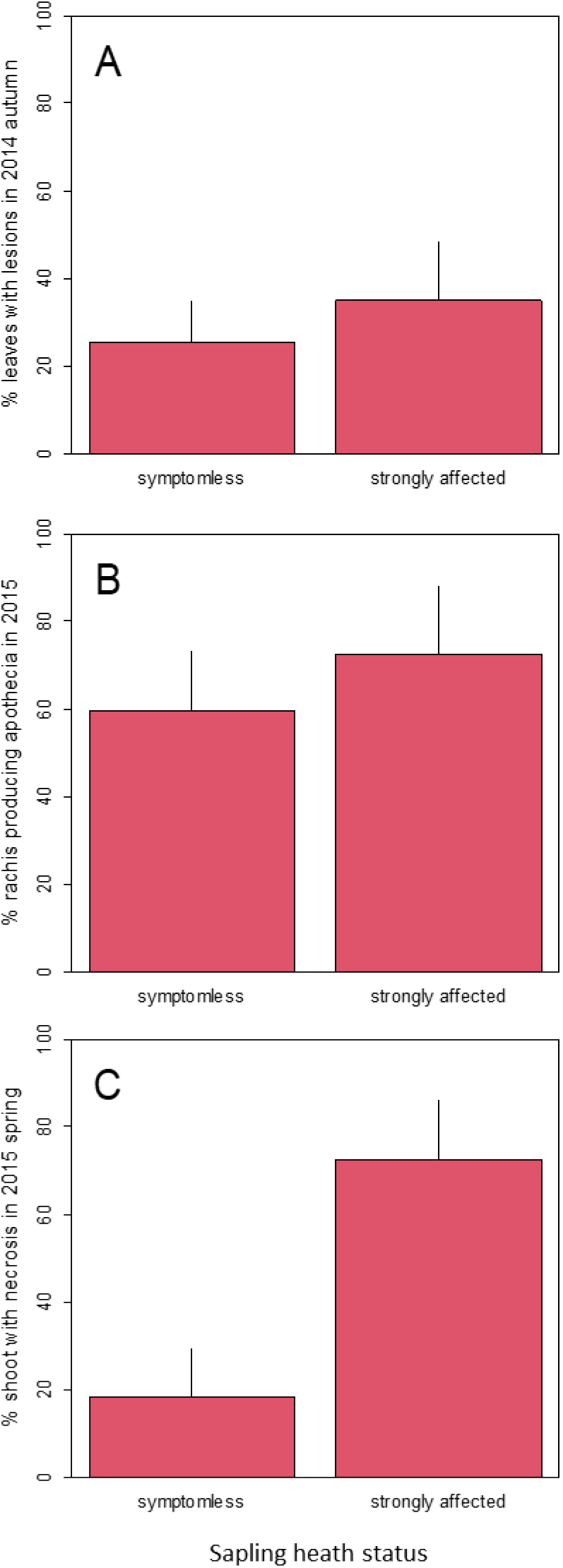
Impact and ability of *H. fraxineus* to complete it cycle on asymptomatic and strongly affected ash seedlings. A. % leaves with necrosis in autumn 2014. B. % of rachis length with presence of *H. fraxineus* apothecia in April 2015, C. % of rated shoot with bark infection in summer 2015.

*H. fraxineus* was able to produce apothecia on rachis of both the asymptomatic and strongly affected saplings at the same rate (likelihood Chi-square = 1.19, p= 0.276, Fig 2b). In summer 2015, the general heath of the saplings had not changed, i.e. only very minor sign of *H. fraxineus* infection was observed on asymptomatic saplings. The likelihood of shoot infection was significantly higher in strongly affected than in asymptomatic saplings (likelihood Chi-square = 6.99, p= 0.008, Fig 2c).

### Relationship between leaf necrosis in fall and subsequent shoot mortality

Shoot mortality in spring was strongly related to leaf necrosis likelihood at the end of the previous summer (Fig. 3a, b, Table 1). It is noteworthy that shoots that showed no leaf necrosis in September had nevertheless a small likelihood to die in the next spring (Fig 3a). Shoot mortality was very different on ashes with different health status (Table 1, Fig 3a). The odds ratio for shoot mortality of a declining ash versus a healthy one was 4.1 (credible interval CI95% [2.4, 7.0]). By contrast, leaf necrosis did not significantly depend on tree health status (Table 1, mean leaf necrosis frequency of 12.9 ± 3.8 % for healthy ashes and 16.3 ± 6.3 % for declining ashes). Both leaf necrosis and shoot mortality were strongly affected by the tree random factor with a similar range of standard deviation (Table 1). Some trees consistently showed more leaf necrosis across years.

**Fig. 3.**
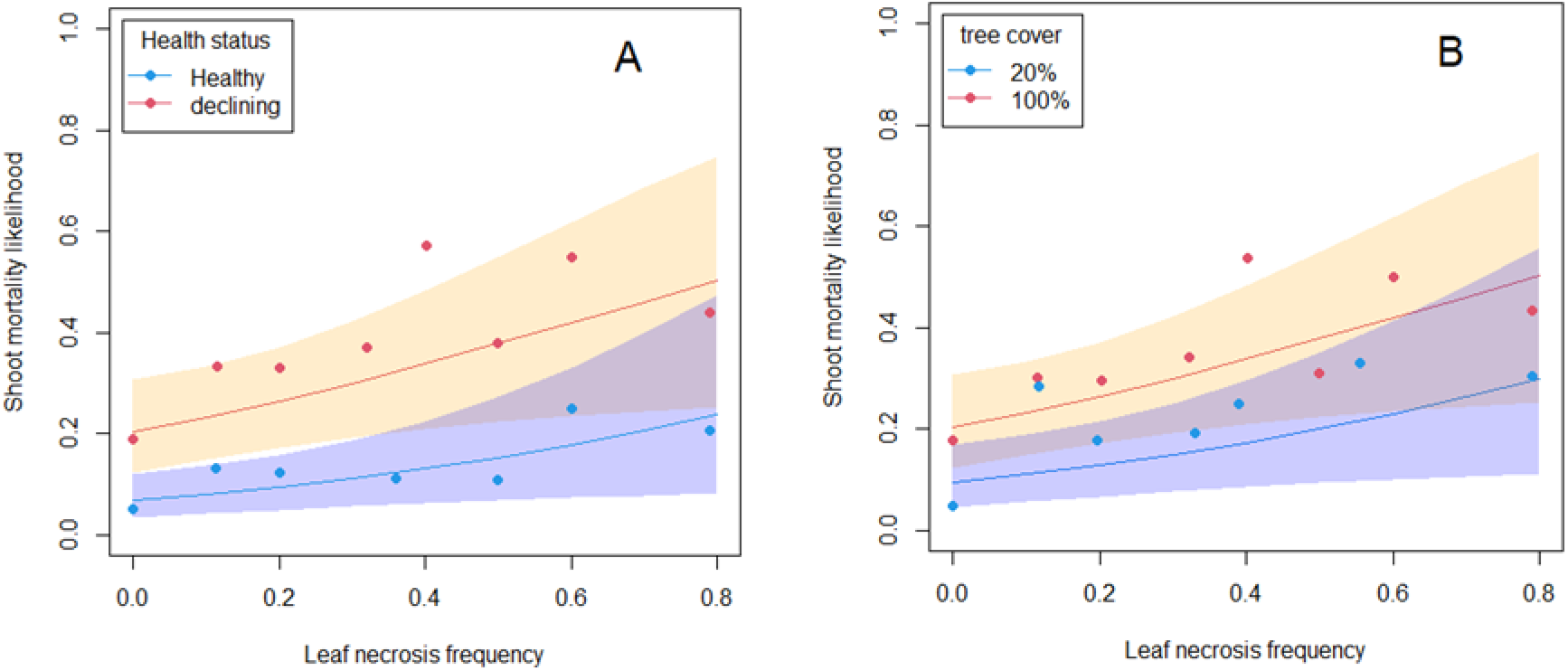
Modelled and observed shoot mortality for Ash trees of different heath status (A) of in sites with different tree cover (B). Point represent the observed data, pooled by increasing 10% leaf necrosis frequency. For tree cover, blue dot represent sites with 9-25 tree cover while red dot represent sites with 75-100 % tree cover. Points representing less than 10 shoots are not represented.

**Table 1.** Relationship between leaf necrosis and shoot mortality at the shoot level. Parameters of the hierarchical Bayesian model

| Variable | Leaf necrosis | Shoot mortality |
| --- | --- | --- |
| Sapling health status (HS) | 0.27 [-0.01, 0.55] | **1.27 [0.82, 1.75]** |
| Site tree cover (TC) | -0.01 [-1.14, 1.16] | **1.40 [0.07, 2.23]** |
| Defoliation at September rating (%DEF) | - | **1.53 [0.38, 2.68]** |
| Probability of necrotic leaves (pl) | - | **1.75 [0.24, 3.27]** |
| Tree random factor (sd, Tree.l and Tree.s) | 0.93 [0.82, 1.06] | 0.89 [0.55, 1.24] |
| Site*year random factor (sd, Env.l and Env.s) | 0.95 [0.65, 1.32] | 1.22 [0.97, 1.54] |
| Deviance | 4615 [4532, 4698] |  |
| NOTE. Parameter [0.95% credible interval]. Parameter significantly different from 0 are marked in bold. |  |  |

Shoot mortality was also significantly affected by tree cover (Table 1, Fig 3b), with an odds ratio of 1.2 (IC [1.0, 1.3]) per 10% tree cover. By contrast, leaf necrosis was not affected by tree cover (Table 1). The environment random factor induced a level of variability similar for leaf necrosis and shoot mortality (Table 1).

Early defoliation was significantly associated with shoot mortality, with increasing shoot mortality with increasing defoliation level (Table 1). Defoliation appeared to be a consequence of infection by *H. fraxineus* and significantly increased with leaf necrosis likelihood (likelihood Chisq of 58.3, p< 0.0001).

### Relationship between leaf necrosis or shoot mortality and climate

The preliminary analysis showed that the best fit for foliar necrosis occurred for climatic conditions in June although the difference was minor (BIC of −103 compared to BIC of −102 for 15 May to 15 June and −97 for 15 June to 15 July). Likewise, for shoot mortality, the best fit occurred for climatic conditions in September to end of December (BIC of 223 compared to BIC of 233 for climatic conditions in September-October).

The model fitted adequately the data, with a proper residual distribution and an acceptable prediction of observed values in the cross-validation (Fig. S1). The main parameter influencing leaf necrosis was TX78, the average of daily maximal temperature in July and August with a strong negative effect of high summer temperatures (Table 2, Fig. 4a). High average temperature in June increased the likelihood of leaf necrosis (Table 2). By contrast, rain either in June or in July-August was not related to leaf necrosis likelihood. Lastly, we observed a trend of decreasing leaf necrosis likelihood with the time of Ash dieback in the area (Table 2, Fig. 4c).

**Table 2.** Relationship between leaf necrosis or shoot mortality and climate parameter (plot level). Parameters of the hierarchical Bayesian model

| Variable | Symptom | Parameter |
| --- | --- | --- |
| Rain June (RA6, $\alpha_2$) | Leaf necrosis | 0.04 [-0.16, 0.25] |
| Temperature June (TM6, $\alpha_3$) | Leaf necrosis] | **0.69 [0.37, 0.99]** |
| Rain Junly-August (RA78 ($\alpha_4$) | Leaf necrosis | -0.20 [-0.52, 0.11] |
| Average daily maximal temperature July-August (TX78, $\alpha_5$) | Leaf necrosis | **-1.11 [-1.50, -0.69]** |
| N year ash dieback presence (LDP, $\alpha_6$) | Leaf necrosis | **-0.37 [-0.58, -0.17]** |
| Site random factor (Site.l, sd) | Leaf necrosis | 0.21 [0.01, 0.49] |
| Rain September-December (RA912, $\beta_2$) | Shoot mortality | -0.06 [-0.43, 0.29] |
| Temperature September-December (TM912, $\beta_3$) | Shoot mortality | **-0.83 [-1.30, -0.39]** |
| Site tree cover (TC, $\beta_4$) | Shoot mortality | **0.77 [0.07, 1.52]** |
| Probability of necrotic leaves (pl, $\beta_5$) | Shoot mortality | **4.55 [4.38, 8.12]** |
| Site random factor (Site.s, sd) | Shoot mortality | 1.21 [0.68, 2.01] |
| Deviance | 3745 [3667, 3826] |  |

NOTE. Parameter [0.95% credible interval]. Parameter significantly different from 0 are marked in bold. The variables are standardized to have zero mean and one standard deviation with the exception of pl, the leaf necrosis likelihood.

**Fig. 4.**
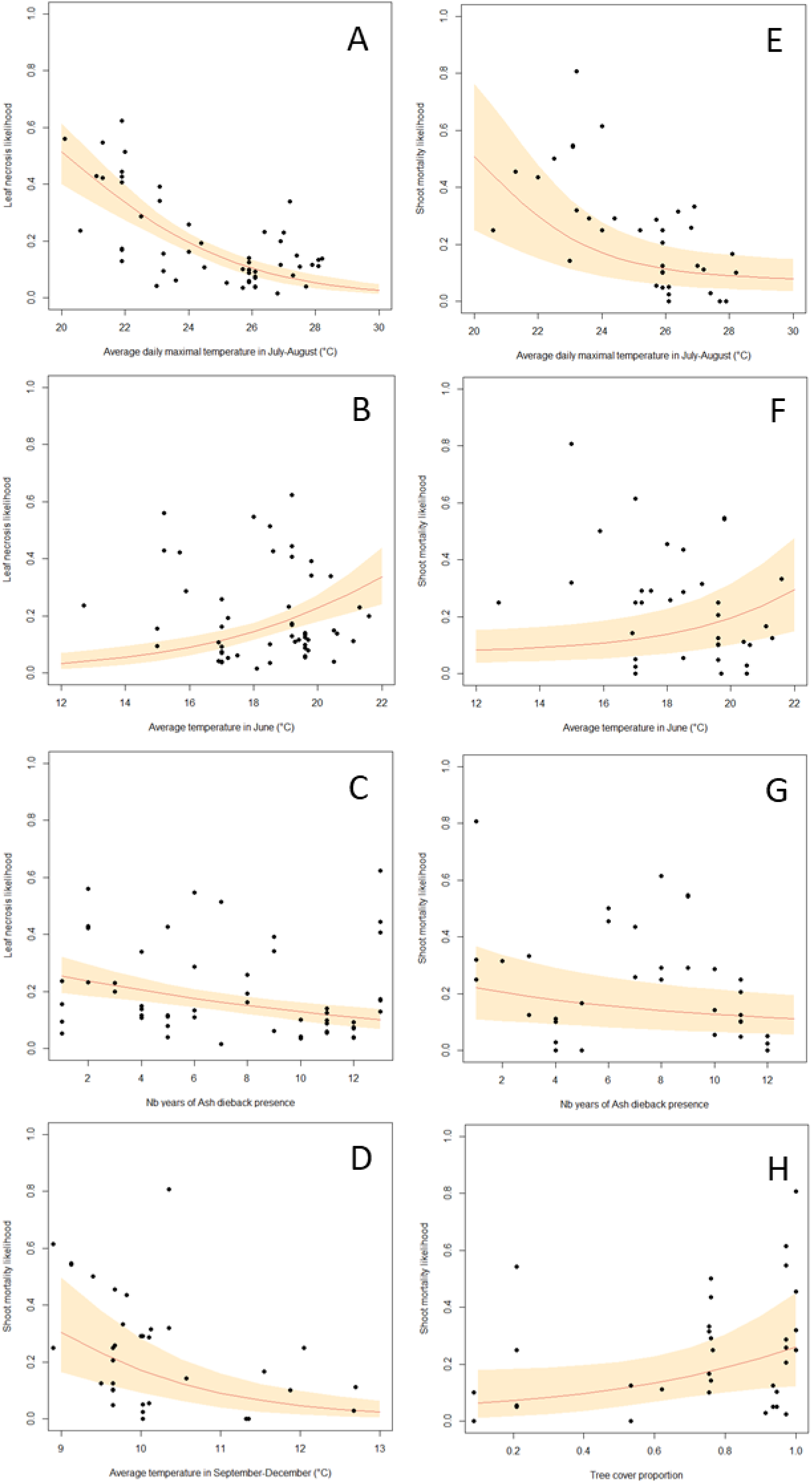
Modelled and observed leaf necrosis and shoot mortality depending on climatic parameters and sites conditions. The curves represent the predicted response for a given parameter, all other parameters being fixed at the average value. The orange area (grey in printed version) represent the 95% confidence interval. The dot represent oberved values.

The shoot mortality was mainly linked to the average temperature in autumn (September to December, Table 2, Fig. 4d) with a decreasing mortality likelihood with increasing autumn temperatures. Rain in autumn was not related to shoot mortality (Table 2). As in the shoot level model, shoot mortality was strongly linked to leaf necrosis likelihood and increased with increasing tree cover (Table 2, Fig. 4g). As a consequence, all parameter influencing leaf necrosis likelihood also influenced shoot mortality (Fig. 4 e, f and g).

### Ability of *H. fraxineus* to produce apothecia on overwintered rachises in the fructification assay

The colonisation of rachises at leaf fall in 2020 and 2021 was tested in part of the sites. It showed striking differences between years and locations (Table 3). Moreover, it was not significantly linked to the leaf necrosis frequency measured in September, two months before (Pearson R= 0.208, pvalue= 0.440). In Champenoux, although the frequency of leaf necrosis was low both in 2019 and 2020, the proportion of rachis in which *H. fraxineus* was detected was very different between the two years (Table 3). Another striking feature was that the frequency of *H. fraxineus* presence in rachises at leaf fall was almost ten-fold higher than the frequency of necrotic leaves.

**Table 3.** Colonisation of rachises at leaf fall in November

|  | Year | Frequency of necrotic leaves in September | Frequency of *H. fraxineus* detection on rachises in November |
| --- | --- | --- | --- |
| Champhenoux | 2019 | 0.09 ± 0.03 | 0.87 ± 0.19 |
|  | 2020 | 0.05 ± 0.02 | 0.35 ± 0.12 |
| Pyrenean piedmont | 2020 | 0.12 ± 0.05 | 0.60 ± 0.18 |

The proportion of rachises producing apothecia in the fructification assay in laboratory conditions showed very high variability, with a range of 0.01 to 0.99. This strongly depended on the year with mean values of 0.26 ± 0.11 in 2016, 0.52 ± 0.17 in 2017, 0.79 ± 0.09 in 2019 and 0.37 ± 0.23 in 2020. The proportion of rachises producing apothecia was significantly correlated to the proportion of rachises in which *H. fraxineus* was detected by qPCR at leaf fall (Pearson R= 0.710, pvalue= 0.002).

The proportion of rachises producing apothecia in the fructification assay was not significantly related to the frequency of leaf necrosis in the previous September rating (Fig. 5a). It was not significantly related to the tested climatic parameters during the overwintering period (Table 4). By contrast, it was significantly related to the rain during the previous summer (sum of rain in July-August, Table 4, Fig. 5b). It was also not related to the % of tree cover (Table 4).

**Fig. 5.**
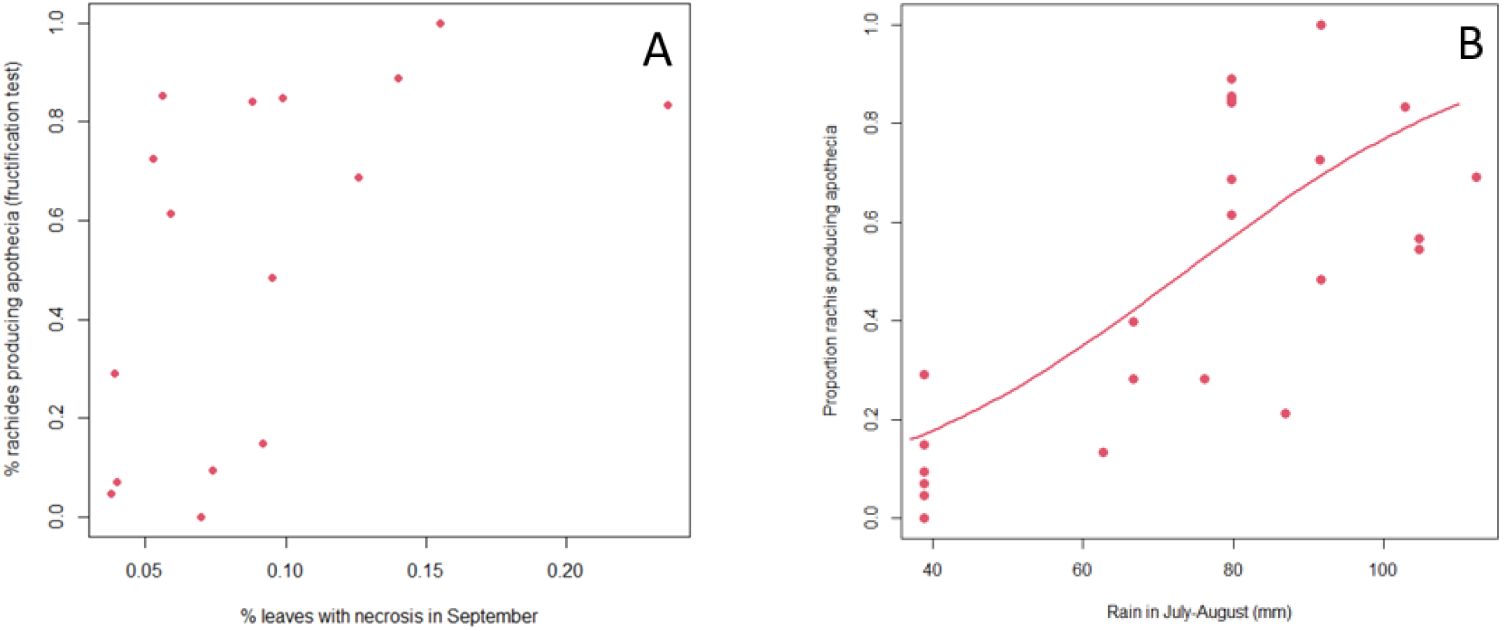
Ability of *H. fraxineus* to produce apothecia on the ash rachises overwintered in common garden (fructification assay). A. relation with leaf necrosis observed in the same plot two months before (pvalue= 0.304); B. relation with climate (pvalue= 0.006, Table 4).

**Table 4.** Influence of climatic parameters on the ability of *H. fraxineus* to produce apothecia in the fructification assay

| Variable | Parameter | Pvalue |
| --- | --- | --- |
| Leaf necrosis frequency | -4.79 | 0.304 |
| Rain in July-August (RA78) | 1.41 | 0.006 |
| Mean maximal daily temperature in July-August (TX78) | -0.32 | 0.411 |
| Rain in October-March (RA10-3 ow) | 0.26 | 0.613 |
| Mean temperature in October-March (TM10-3 ow) | 0.46 | 0.112 |
| Site tree cover (% TC) | 0.43 | 0.444 |
| Site random factor (sd) | 0.31 | - |

Note: RA10-3 ow and TM10-3 ow, sum of rain and mean of daily temperatures at the overwintering sites from October to March.

### Climate suitability in France for the different stage of *H. fraxineus* life cycle

Maps were produced using the developed models (Table 2 and 4) for an average site (site random factor fixed at 0) and for years 2010 to 2020. *H. fraxineus* should be able to colonise leaves, be present on rachises shed in autumn and be able to produce apothecia in spring in most of France (Fig 6a). By contrast, its ability to produce damage should be much more limited, in particular in southern France (Fig. 6b, c and d). However, its ability to cause damage to ash trees is strongly dependent on local site conditions: comparison of Fig 6c and Fig 6d show large difference of prediction shoot mortality likelihood depending on the % tree cover. Uncertainty in parameter estimation very significantly affected both the maps of necrotic leaves / shoot mortality probability (Fig. S2).

**Fig. 6.**
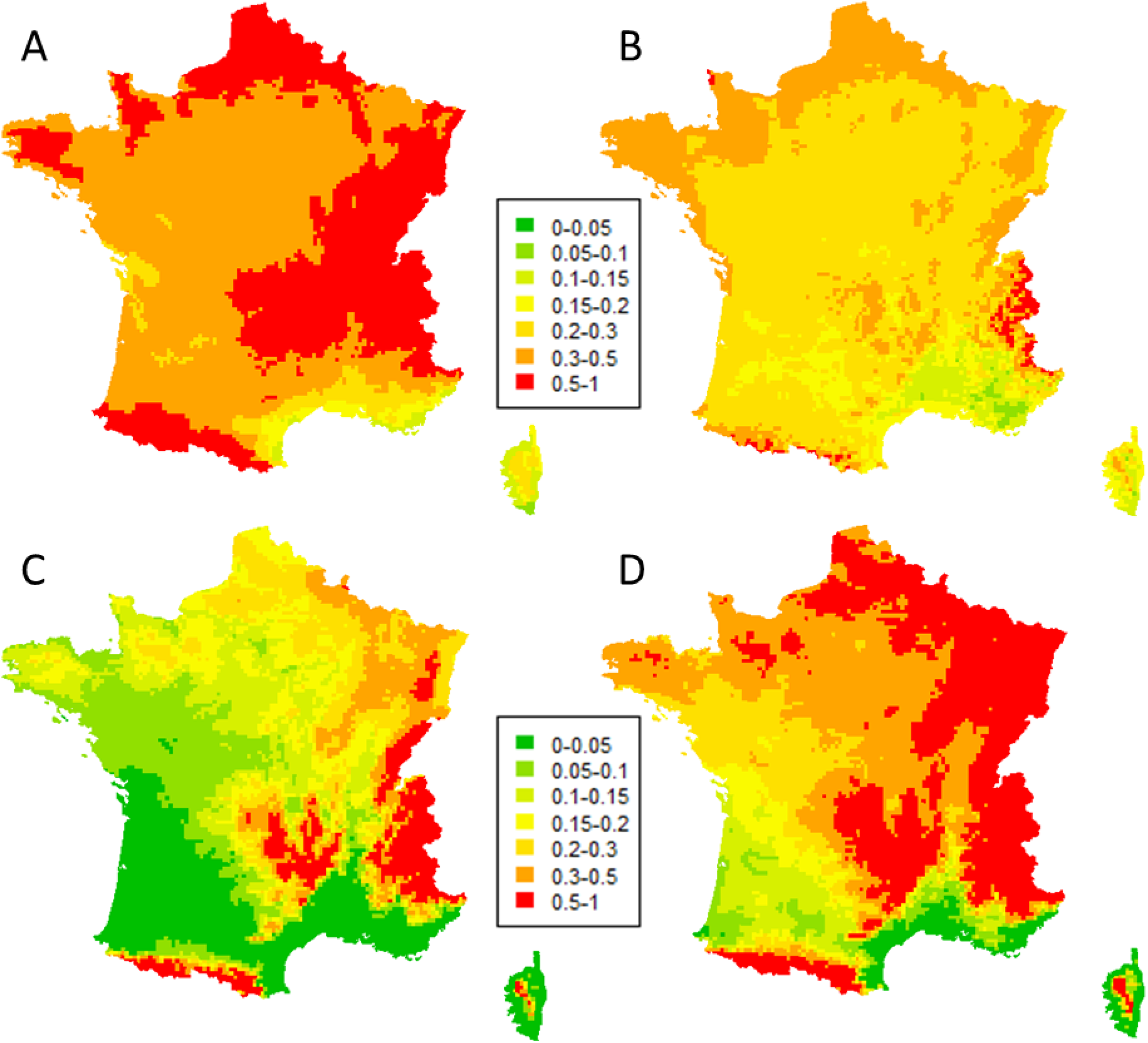
Map of climate suitability for *H. fraxineus* ability to produce apothecia on the ash rachises or symptoms on leaf and shoot of Ash trees (Safran meteorological data, 2010-20). A. Proportion of Ash rachises collected in fall producing apothecia in laboratory conditions after overwintering in common garden (fructification assay); B. Leaf necrosis likelihood; C. Shoot mortality likelihood in open canopy (20% tree cover); D Shoot mortality likelihood in forest conditions (100% tree cover).

## Discussion

We showed that healthy carrier exists and play a significant role in ash dieback epidemiology. *H. fraxineus* is able to establish on ash leaves and to reproduce on the leaf debris in the litter (rachises) in situations, either local environment, meso-climates or specific trees where it induce very limited shoot mortality. Climate strongly influence *H. fraxineus* with different parameters being important depending on the life cycle stage. As a consequence, while the pathogen might be able to establish over most France, it should not be able to induce significant dieback in large part of southern France. Last, we observed a decreasing trend of severity (leaf necrosis and shoot mortality likelihood) with the time of disease presence.

We were able to predict the severity of dieback in spring by observing leaf symptoms in the previous autumn. Leaf necrosis frequency in September was a very good predictor of subsequent shoot mortality. Despite that, leaf necrosis was a very poor predictor of rachises infection at leaf fall and of the ability of *H. fraxineus* to produce apothecia, and therefore inoculum on the rachises in spring. This may be because of the two month delay between leaf necrosis measurement and rachises colonisation assessment. It has to be pointed out that very little inoculum production is observed September and October in French conditions (Grosdidier et al, 2018). Nevertheless, lesions present in the leaflets might have extended in the rachises. Another likely explanation is that *H. fraxineus* has the ability to asymptomatically infect ash leaves in early summer and to behave as a latent pathogen for an extended period (Cross et al, 2017). These authors showed that leaf necrosis occurs only when *H. fraxineus* inoculum reaches a certain level in the leaf which happens at the same time as a change in the leaf microbiota composition. It would appear that the high *H. fraxineus* levels in leaves required for leaf necrosis also strongly promote shoots infection from the petiole. By contrast, it may not be required for rachises colonisation at leaf fall in senescent tissue; this would explain the observed result.

As a consequence, *H. fraxineus* is able to complete its biological cycle in conditions where no significant dieback is observed and some ashes behave as healthy carriers. This is the case at local level, on specific trees that retain healthy crown despite experiencing leaf necrosis and enabling the pathogen to complete its cycle on infected rachises. Our experimental design did not allow to determine whether these trees remains heathy for genetic reasons, because they were in microsite unfavorable to the disease or because foliar microbial competitors prevent *H. fraxineus* development. Resistance toward *H. fraxineus* has been widely documented in *F. excelsior* (McKinney *et a*l, 2011, Semizer-Cuming *et al*, 2021) and thus remains a possible explanation for at least part of the observed pattern. It does not however appear that the studied trees remained healthy because they avoided inoculum as they showed a frequency of leaf necrosis similar to trees that showed dieback symptoms in the same plots. In the two stands studied in more detail in 2014, the ash density was high and the saplings remaining healthy were in close proximity to dying saplings, questioning the possibility that they were experiencing different environmental conditions. Nevertheless, we do have evidence that environmental conditions strongly influenced the ability of *H. fraxineus* to infect the shoot from the leaf petiole. Indeed, although the likelihood of leaf necrosis did not depend on the plot tree cover, the subsequent likelihood of shoot mortality was much lower in hedges with a low tree cover compared to forest plots with a high tree cover. We also showed that the colonisation of leaf rachises at leaf fall and subsequent ability of *H. fraxineus* to complete its cycle on the colonised rachises does not depend on the tree cover, being similar in forests conditions and in hedges in open landscape. Hence, trees with no dieback symptoms in infected stands behave as heathy carriers, allowing *H. fraxineus* to reproduce on their leaves as efficiently as on the leaves of neighboring trees that experience dieback. They should even be better inoculum producers than trees affected by dieback as they retain full foliage and thus produce abundant infected rachises. This result is in agreement with the observation of Grosdidier et al (2020) that the density of infected rachises was similar in the litter beneath ash trees in hedges and in forest settings although the trees showed far less dieback symptoms in the latter situation. A possible explanation is that as the percent tree cover does not affect inoculum production, it does not affect infection frequency. However, at low tree cover, crowns are exposed more frequently to temperature adverse to *H. fraxineus* (Grosdidier & al, 2020), preventing the increase of the pathogen present as endophyte in leaves and thus later shoot mortality. This discrepancy between the ability to reproduce on the host and the ability to induce significant damages has been reported for other tree pathogens. For example, *Diplodia sapinea* mainly produce inoculum on Pine cones and is widely present on cones in healthy Pine stands; it however needs weakening of the trees to induce significant damages (Palmer et al, 1988, Blodgett et al, 1997).

At a larger scale, we showed that *H. fraxineus* may be able to fulfil its cycle in regions were climate does not allow significant dieback. The disease has been progressing very regularly in France from 2009 to 2016 with a very noticeable slowdown of the spread in southern France from 2017 to 2019 (Fig 1b). The slowdown of the spread occurred in regions with climate unfavorable to ash dieback. In 2020, the Pyrenean area was colonised while no sign of the disease had been reported north in the Garonne valley (DSF database). It is possible that the pathogen was introduced via infected planted ashes (Wylder *et al*, 2018). However, our results show that the spread without dieback symptoms of *H. fraxineus* in the Garonne valley is also a likely hypothesis. This may have been enhanced by the very hot summers of 2018-20, which were even less favorable to *H. fraxineus*-induced shoot mortality. The delay of 3-4 years observed until the first report of ash dieback in the Pyrenean piedmont is in line with the spread speed observed in France of about 60 km per year (Grosdidier, 2017).

The decreasing trend of leaf necrosis and shoot mortality likelihood with years of disease presence may be very significant but requires confirmation. It may well be a spurious result caused by the very dry and hot climate experienced in northern and eastern France in 2018-20, at the end of the study period. Climatic conditions were taken into account in our model and summer high temperature was one of the most important feature to explain symptoms severity. But the parameter we used (average daily maximal temperature) may not have accounted to entire effect of heat waves observed during this 2018-20 period. Another possible explanation is the progressive selection of more tolerant ashes in sites affected by ash dieback since a long time. Indeed, in the site followed for the longest period, less than 50% of the seedlings selected in 2015 were still alive in 2021. We tried to replace dead saplings by symptomatic saplings to avoid a progressive increase in tolerant ash frequency in our sample, but this was not always possible. Alternatively, other environmental features might have changed such as the microbiota associated with leaves or rachises in the litter (Desprez-Loustau & al, 2019).

Early defoliation was hypothesized to be a mechanism of tolerance to *H. fraxineus*, the pathogen then being shed in the leaves before getting the chance to infect shoot (Mc Kinney *et al*, 2011). We were not able to confirm the hypothesis, early defoliation increasing shoot mortality in our data. However, cares have to be taken when interpreting this result as Mc Kinney *et al* (2011) have reported early defoliation as a genetically controlled trait that differed between ashes clones while, in our data, early defoliation was strongly driven by environmental conditions, being tightly associated with strong leaf necrosis frequency. This may have changed the relationship.

Climate was a major driver of symptoms induction by *H. fraxineus*. It was expected that high summer temperatures were unfavorable to the disease, as the pathogen has been shown to have low survival at temperature above 35°C and to be limited in south-east France by the hot summers of the Rhône valley south of Lyon (Hauptman *et al*, 2013, Grosdidier *et al*, 2018). More surprising is the low influence of spring and summer rains as those have been associated with high disease impact through increased inoculum production (Dvořák et al, 2016, Skovsgaard *et al*, 2017, Chumanová *et al*, 2019). Spring and summer rain may promote infection of the leaves, but *H. fraxineus* can then remain as asymptomatic endophyte especially if high summer temperatures prevent the heavy colonisation of ash leaves by the pathogen. This could explain why spring and summer precipitations were not well correlated to the induction of symptoms such as leaf necrosis and shoot mortality. By contrast, summer rainfall was the main driver of rachises infection and of their ability to produce apothecia in the following spring. This is the likely mechanism by which spring and summer rainfall favour high ash dieback impact. The period in spring with the best fit between climatic condition and leaf necrosis likelihood was in June. This may appear surprising as the period of ascospores production has often been shown to be in summer, in July to August (Chandelier *et al*., 2014; Dvorak *et al*., 2016; Hietala *et al*., 2013; Timmermann *et al*., 2011). However, many of those results correspond to central or northern Europe; in NE France, the period of ascospores production was earlier, from mid-June to mid-July and ascomata have been reported as early as May in SW France (Grosdidier *et al*., 2018, C. Husson, unpublished result). Shoot mortality was associated with previous leaf necrosis, but also to subsequent mild autumn temperature. This association of shoot mortality with autumn climatic conditions was by now never reported in the literature and the mechanism involved remains to be elucidated.

*H. fraxineus* ability to infect ash leaves and reproduce on the rachises in the forest litter does not appears to limit the pathogen ability to establish throughout France. By contrast, the ability to induce shoot mortality is the limiting step. Hence, the maps showing the shoot mortality likelihood match well with the ash dieback risk predicted by Goberville *et al* (2016). The statistical niche model produced by these authors was based on the distribution in Europe of this invasive pathogen still spreading and with limited presence in France; it nevertheless well captured the potential climatic niche in France. This would confirm that south-eastern France is not favorable to ash dieback, as was suggested by Grosdidier et al (2018).

## Acknowledgement

We wish to thanks Olivier Caёl for technical assistance in handling the analysis of rachises ability to produce apothecia and level of infection by *H. fraxineus*. The establishment and rating of the plots in the Charente area was done by Dominique Piou while those of plots from Brittany was done by Laurence Roche from the DSF. We also thank Mireia Gomez-Gallego for her advices in the analysis and for reviewing the manuscript. This work was funded by the French Forest Health Department, the French Ministry of Agriculture and Forestry and by the H2020 EU contract HOMED. The UMR1136 research unit is supported by a grant managed by the French National Research Agency (ANR) as part of the *“Investissements d’Avenir*” program (ANR-11-LABX-0002-01, Laboratory of Excellence ARBRE).

## Supplemental data

**Table S1.** Studied sites

| Plot | Name (Region) | Coordinates | Leaf infection | Shoot mortality | % tree cover |
| --- | --- | --- | --- | --- | --- |
| Am | Fréchencourt (Haut-de-France) | 2.43261 ; 49.97479 | 2016-20 | 2017-21 | 76 |
| Ch1 | Champenoux (Grand-Est) | 6.34521 ; 48.75327 | 2015-21 | 2016-22 | 97 |
| Ch2 | Champenoux (Grand-Est) | 6.31782 ; 48.73877 | 2016-21 | 2017-22 | 21 |
| Ch3 | Champenoux (Grand-Est) | 6.33035 ; 48.75654 | 2019-21 | 2020-22 | 95 |
| Ch4 | Champenoux (Grand-Est) | 6.35984 ; 48.73698 | 2019-21 | 2020-22 | 3 |
| Ch5 | Champenoux (Grand-Est) | 6.37312 ; 48.74222 | 2019-21 | 2020-22 | 94 |
| Ch6 | Champenoux (Grand-Est) | 6.32135 ; 48.73938 | 2019-21 | 2020-22 | 9 |
| Gr | Gremecey (Grand-Est) | 6.43011 ; 48.81474 | 2014-15 | 2015 | 100 |
| Se | Seichamps (Grand-Est) | 6.26910 ; 48.72059 | 2013-14 | <sup>a</sup> | 15 |
| Lu | Lupé (Auvergne-Rhône-Alpes) | 4.71034 ; 45.37561 | 2016-20 | 2017-20 <sup>a</sup> | 75 |
| Ro | Roche-sur-Grane (Auvergne-Rhône-Alpes) | 4.93764 ; 44.68831 | 2016-20 | 2017-20 <sup>b</sup> | 63 |
| Co | Colombier (Nouvelle-Aquitaine) | -0.55205 ; 45.66159 | 2018-19 | 2019 <sup>a</sup> | 62 |
| Sa | Salignac-sur-Charente (Nouvelle-Aquitaine) | -0.39385 ; 45.66485 | 2018-19 | 2019 <sup>a</sup> | 91 |
| Be | Sarrance (Nouvelle-Aquitaine) | -0.58197 ; 43.02852 | 2020 | 2021 | 77 |
| Ol1 | Oloron-Sainte-Marie (Nouvelle-Aquitaine) | -0.54005 ; 43.11422 | 2020 | 2021 | 100 |
| Ol2 | Oloron-Sainte-Marie (Nouvelle-Aquitaine) | -0.53388 ; 43.12531 | 2020 | 2021 | 100 |
| Fe | Ance Féas (Nouvelle-Aquitaine) | -0.65832 ; 43.15170 | 2020 | 2021 | 100 |
| La | Landivisiau (Bretagne) | -4.08828 ; 48.52195 | 2020 | <sup>b</sup> | 42 |
| Po | Pont-de-Buis Les Quimerch (Bretagne) | -4.07908 ; 48.24803 | 2020 | <sup>b</sup> | 95 |
| Sc | Scaër (Bretagne) | -3.70625 ; 48.03131 | 2020 | <sup>b</sup> | 83 |

<sup>a</sup> Data not available for shoot mortality because the site was destroyed during the winter
<sup>b</sup> Data not recorded in spring 2021

**Fig S1.**
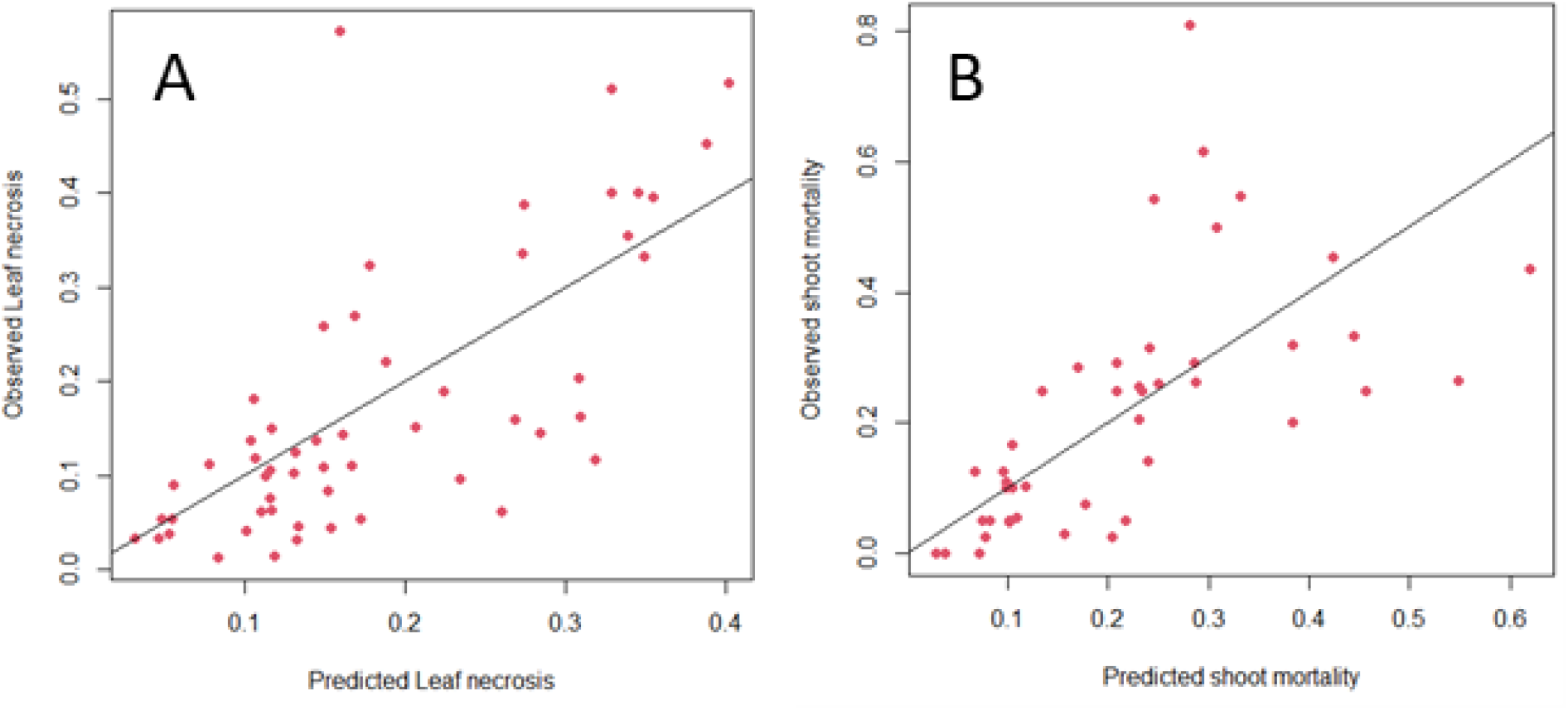
Cross-validation for the model at the plot level relating climatic and site parameters to leak necrosis and shoot mortality likelihood. A. Leaf necrosis, B. shoot mortality

**Fig S2.**
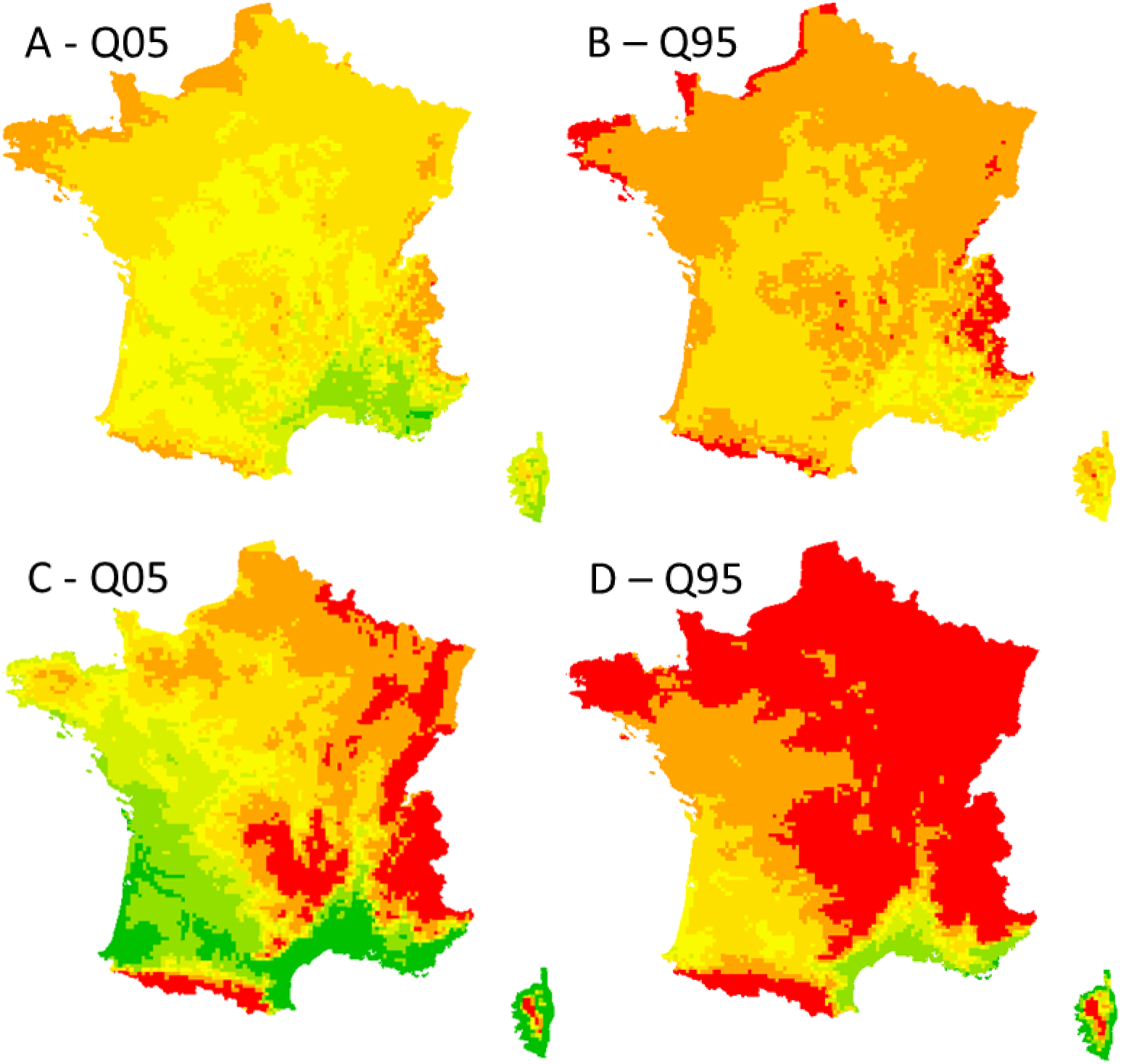
Quantile 0.05 (Q5) and 0.95 (Q95) for the estimated likelihood of leaf necrosis (A, B) and shoot mortality (C, D). The map were computed using the parameter 0.05 and 0.95 quantiles obtained from the Bayesian procedure fit. The shoot mortality is computed for a forest situation (tree cover of 100%).

